# Semantic Integration of Clinical Laboratory Tests from Electronic Health Records for Deep Phenotyping and Biomarker Discovery

**DOI:** 10.1101/519231

**Authors:** Xingmin Aaron Zhang, Amy Yates, Nicole Vasilevsky, JP Gourdine, Leigh C. Carmody, Daniel Danis, Marcin P. Joachimiak, Vida Ravanmehr, Emily R. Pfaff, James Champion, Kimberly Robasky, Hao Xu, Karamarie Fecho, Nephi A. Walton, Richard Zhu, Justin Ramsdill, Chris Mungall, Sebastian Köhler, Melissa A. Haendel, Clem McDonald, Daniel J. Vreeman, David B. Peden, Christopher G. Chute, Peter N. Robinson

## Abstract

Electronic Health Record (EHR) systems typically define laboratory test results using the Laboratory Observation Identifier Names and Codes (LOINC) and can transmit them using Fast Healthcare Interoperability Resource (FHIR) standards. LOINC has not yet been semantically integrated with computational resources for phenotype analysis. Here, we provide a method for mapping LOINC-encoded laboratory test results transmitted in FHIR standards to the Human Phenotype Ontology (HPO) terms. We annotated the medical implications of 2421 commonly used laboratory tests with HPO terms. Using these annotations, a software assesses laboratory test results and converts each into an HPO term. We validated our approach with EHR data from 15,681 patients with respiratory complaints and identified known biomarkers for asthma. Finally, we provide a freely available SMART on FHIR application that can be used within EHR systems. Our approach allows reusing readily available laboratory tests in EHR for deep phenotyping and using the hierarchical structure of HPO for association studies with medical outcomes and genomics.

**One Sentence Summary:** We present an approach to semantically integrating LOINC-encoded laboratory data with the Human Phenotype Ontology and show that the integrated LOINC data can be used to identify biomarkers for asthma from electronic health record data.

## Introduction

Electronic health records (EHRs) have been widely adopted in US hospitals since the Health Information Technology for Electronic and Clinical Health Act (HITECH) was passed in 2009, and offer an unprecedented opportunity to accelerate translational research because of advantages of scale and cost-efficiency as compared to traditional cohort-based studies^1^. In particular, EHRs contain rich phenotype information that can be utilized to stratify diseases and to develop hypotheses. For instance, phenome-wide association studies (PheWAS) can exploit EHR data to define case control cohorts for disease diagnoses or laboratory traits and then analyze associations with hundreds of thousands of genetic variants^2–4^. Despite the great potential of EHR data, patient phenotyping from EHRs is still challenging because the phenotype information is distributed in many EHR locations (laboratories, notes, problem lists, imaging data, etc.) and with EHRs having vastly different structures across sites. This lack of integration represents a substantial barrier to widespread use of EHR data in translational research.

Laboratory tests provide a critical resource for phenotype extraction. Deep phenotyping, i.e., comprehensive and precise phenotyping of individual disease manifestations, is an essential component of precision medicine and could potentially extend the reach of PheWAS studies^5,6^. Laboratory tests have broad applicability for translational research, but EHR-based research using laboratory data has been challenging because of the lack of standardization among different EHR systems. For instance, some tests measure nitrite level in urine using an automated machine, whereas others use a test strip. Some report the value in mg/dL whereas others report a qualitative value of positive/negative. If any of these tests were abnormal, the medical interpretation would be that nitrituria is present, yet current informatics frameworks do not easily support such inferences. Therefore, substantial challenges exist for standardization and integration of laboratory data for deep phenotyping and EHR-based translational research.

Recent advances in the standardization of EHR systems and phenotyping ontologies make it feasible to extract patient phenotypes from laboratory tests at a large scale. The Fast Healthcare Interoperability Resource (FHIR) was introduced in 2013 and provides a standardized interface to individual EHR systems for healthcare-related data^7^. FHIR separates healthcare-related data into granular components as “resources” such as observation, medication, patient identity and insurance claims, that have a standard definition and associated semantic bindings that can be computationally integrated even when they are created by different methods and organizations. Laboratory tests, encoded as observations in FHIR, are uniquely identified with Laboratory Observation Identifier Names and Codes (LOINC), which is a universal code system that defines various kinds of clinical laboratory tests and other measurements (~86,000 entries)^8^. The outcome of a FHIR observation can be represented by a term in the Human Phenotype Ontology (HPO), which is a logically defined vocabulary for describing human abnormal phenotypes^9^. The HPO has become the de facto standard for computational phenotype analysis in genomics and rare disease^9–11^. The HPO currently contains 13,608 terms including a comprehensive representation of laboratory abnormalities such as *Hyperglycemia, Thrombocytopenia*, and *Increased urine alpha-ketoglutarate concentration*. Here, we present a computational method that semantically harmonizes FHIR, LOINC, and HPO. The software rolls up LOINC terms for tests whose outcomes are medically comparable into common categories and interprets the outcome as HPO terms, thereby automatically extracting detailed, deep phenotypic profiles of laboratory results for downstream studies.

## Results

### Overview of strategy

We present an approach to mapping the outcomes of laboratory tests as encoded in EHRs with LOINC terms for the tests and FHIR Observation resources representing the test results as HPO terms. A LOINC term by itself does not specify the outcome of a test. But if the outcome of a test (such as “high” or “low”) and the nature of the test are known, we can then infer the phenotypic abnormality. For example, LOINC 32710-6 “Nitrite [Presence] in Urine” together with the outcome “positive” implies the phenotypic abnormality *Nitrituria* (HP:0031812).

LOINC-coded laboratory tests can be grouped broadly into three categories, those with a quantitative outcome (Qn), an ordered categorical outcome (ordinal or Ord) and an unordered categorical outcome (nominal, or Nom). A quantitative test for an analyte has a normal range, and there are three types of mappings depending on the result of the test: L (lower than normal), N (normal), and H (higher than normal). Take, for instance, a test for the concentration of potassium in the blood (LOINC:6298-4, Fig. 1A). If the result is high, our procedure infers the corresponding HPO term for *Hyperkalemia* (HP:0002153). Analogously, a low result is mapped to *Hypokalemia* (HP:0002900). The HPO is an ontology of abnormal phenotypes, and thus there is no term that specifically represents a normal test result. However, computational analysis can record negated HPO terms, and the normal test result is represented as NOT *Abnormality of potassium homeostasis* (HP:0011042).

**Fig. 1.**
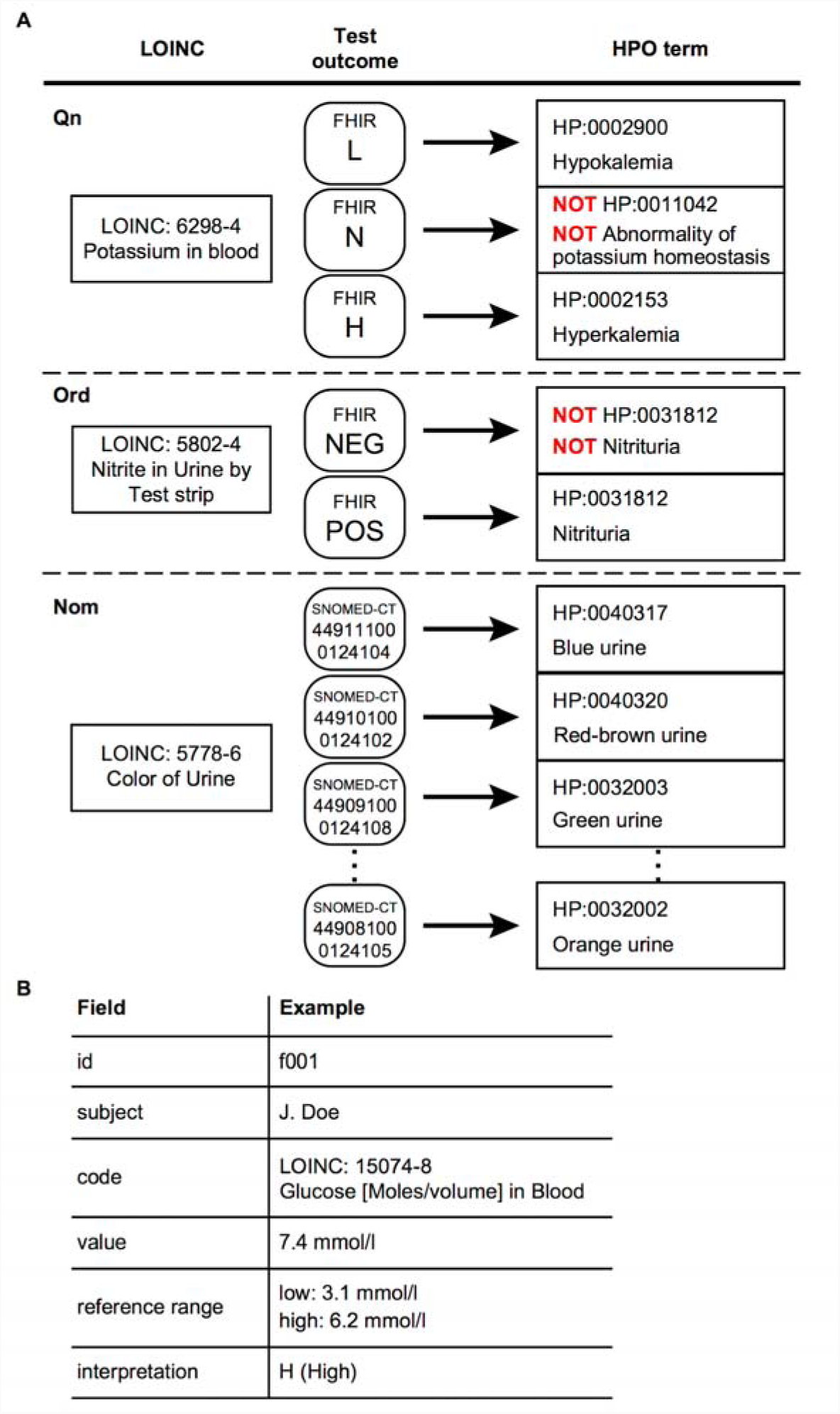
Semantic integration of LOINC-coded laboratory tests in FHIR into HPO terms. **A.** Representative examples of LOINC to HPO mapping. Potassium in blood, a quantitative (Qn) LOINC test, has three potential outcomes, L (lower than normal), N (normal) and H (higher than normal) and is mapped to three corresponding HPO terms. Presence of nitrite in urine, an ordinal (Ord) test, has two possible outcomes, POS (positive) or NEG (negative) and is mapped to either *Nitrituria* (HP:0031812) or NOT *Nitrituria* (HP:0031812), respectively. Color of urine, a nominal (Nom) test, has a list of likely outcomes (represented as Systematized Nomenclature of Medicine-Clinical Terms (SNOMED-CT) codes) and each one is mapped to an HPO term. **B.** Schematic representation of the relevant contents of a FHIR observation for laboratory tests. Each FHIR Observation resource for a LOINC-encoded laboratory test includes an identifier (id) and the name of the patient, the LOINC code and name, the normal reference range and the observed value as well as an interpretation of the result (see Table 1 for a complete list).

Ordinal tests can have a series of ordered outcomes. The majority of the ordinal LOINC tests were mapped to two possible outcomes, POS (positive) or NEG (negative). For instance, the result of the test Nitrite in urine by test strip can be positive (present) or negative (absent) (Fig. 1A). If present, then our approach infers the HPO term *Nitrituria* (HP:0031812); if absent, our approach infers NOT *Nitrituria* (HP:0031812).

**Table 1.** FHIR codes for test outcomes

| Primary Code | Other FHIR codes mapped | Meaning |
| --- | --- | --- |
| A | AA, W | abnormal |
| L | <, D, LL, LU, | lower than normal |
| N | B, I | normal |
| H | >, HH,HU, U, | higher than normal |
| NEG | ND, NR, | not present |
| POS | AC, DET, RR, TOX, WR | present |
| U | HM, IE, IND, MS, NS, null, OBX, QCF, R, S, SDD, SYN-R, SYN-S, VS | unknown |

Nominal tests have a series of outcomes that lack a natural ordering. Yet, some nominal result values are considered abnormal. For instance, LOINC 5778-6, Color of urine. Currently, nine potential abnormal results of this test are mapped to the nine child terms of *Abnormal urinary color* (HP:0012086), including *Red Urine* (HP:0040318) and *Dark urine* (HP:0040319).

### A LOINC to HPO mapping library

We have mapped 2421 LOINC terms to HPO terms. 77.8% of the mapped LOINC tests are Qn, 21.6% Ord and 0.6% Nom (Fig. 2A). Taken together, these LOINC terms mapped to a total of 516 distinct HPO terms. We analyzed the distribution of the number of distinct LOINC terms that were mapped to an individual HPO term. In 56.4% of the cases, two or more LOINC terms are mapped to the same HPO term (mean = 8.5) (Fig. 2B), reflecting the fact that multiple laboratory tests (and associated LOINC terms) have outcomes that we consider to have an equivalent clinical interpretation and can therefore be mapped to the same HPO term.

**Fig. 2.**
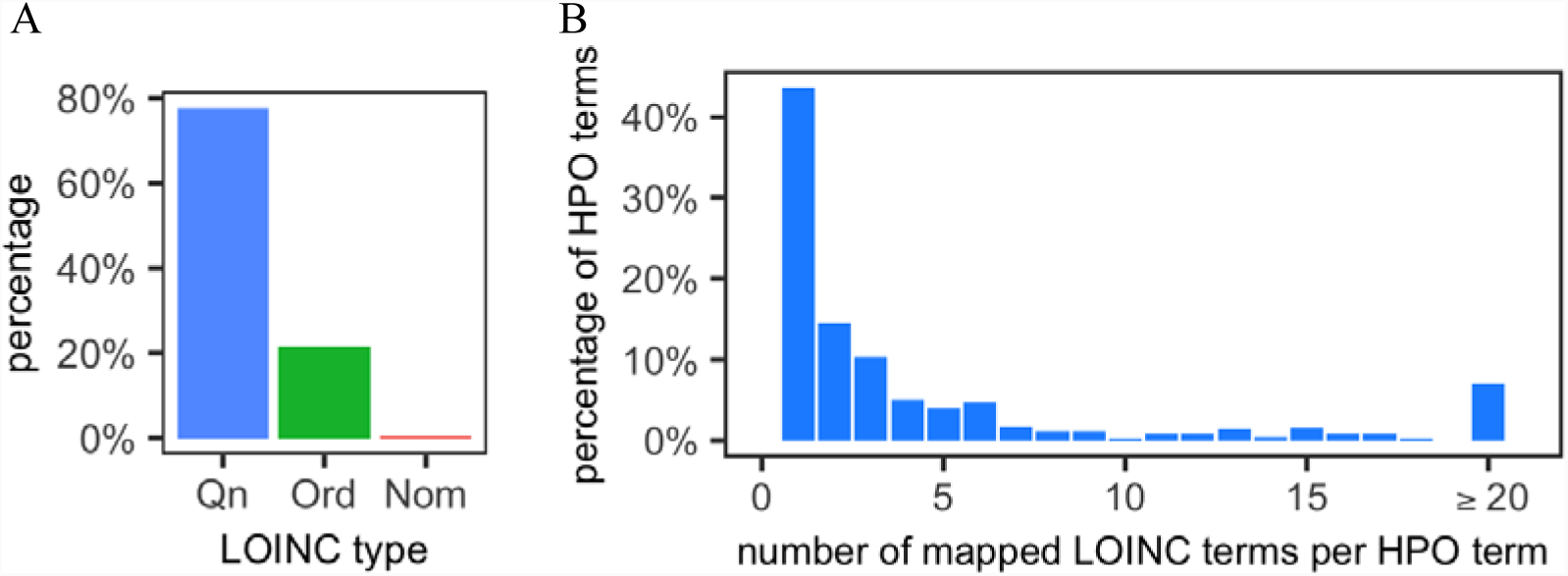
Quantification of the LOINC to HPO mapping library. **A.** Distribution of annotated LOINC terms **B.** Distribution of HPO terms according to the number of LOINC terms mapped to an individual HPO term.

### Algorithm for converting LOINC-coded laboratory tests into HPO-coded phenotypes

We designed an algorithm that inspects elements of a FHIR resource for laboratory tests and converts the outcome into an HPO term. A standard FHIR resource for laboratory tests (a FHIR Observation) contains patient information, test identification, test result, normal reference range and interpretations (Fig. 1B). The algorithm compares the numerical result with the normal reference ranges to assign an interpretation code such as “L” or “POS” (Table 1), or make use of the interpretation codes when they are present, to map the result to the corresponding HPO term (fig. S1). Overall, the algorithm handles all three major types of LOINC-coded laboratory test (Qn, Ord, and Nom) when combined with the LOINC to HPO annotation data.

### HPO on FHIR

To demonstrate conversion of FHIR-encoded LOINC tests into HPO, we created a SMART on FHIR app that uses the mapping library. SMART (Substitutable Medical Applications, Reusable Technologies) on FHIR is an app platform for electronic health records that allows apps to run on different FHIR-enabled EHR systems^12^. Our app, *HPO on FHIR*, transforms a bundle of laboratory observations for a patient into a list of HPO codes (Fig. 3). We have also developed a command-line application that can iterate through all laboratory tests in a FHIR-enabled server, convert each into an HPO term and store them in a relational database for translational research.

**Fig. 3.**
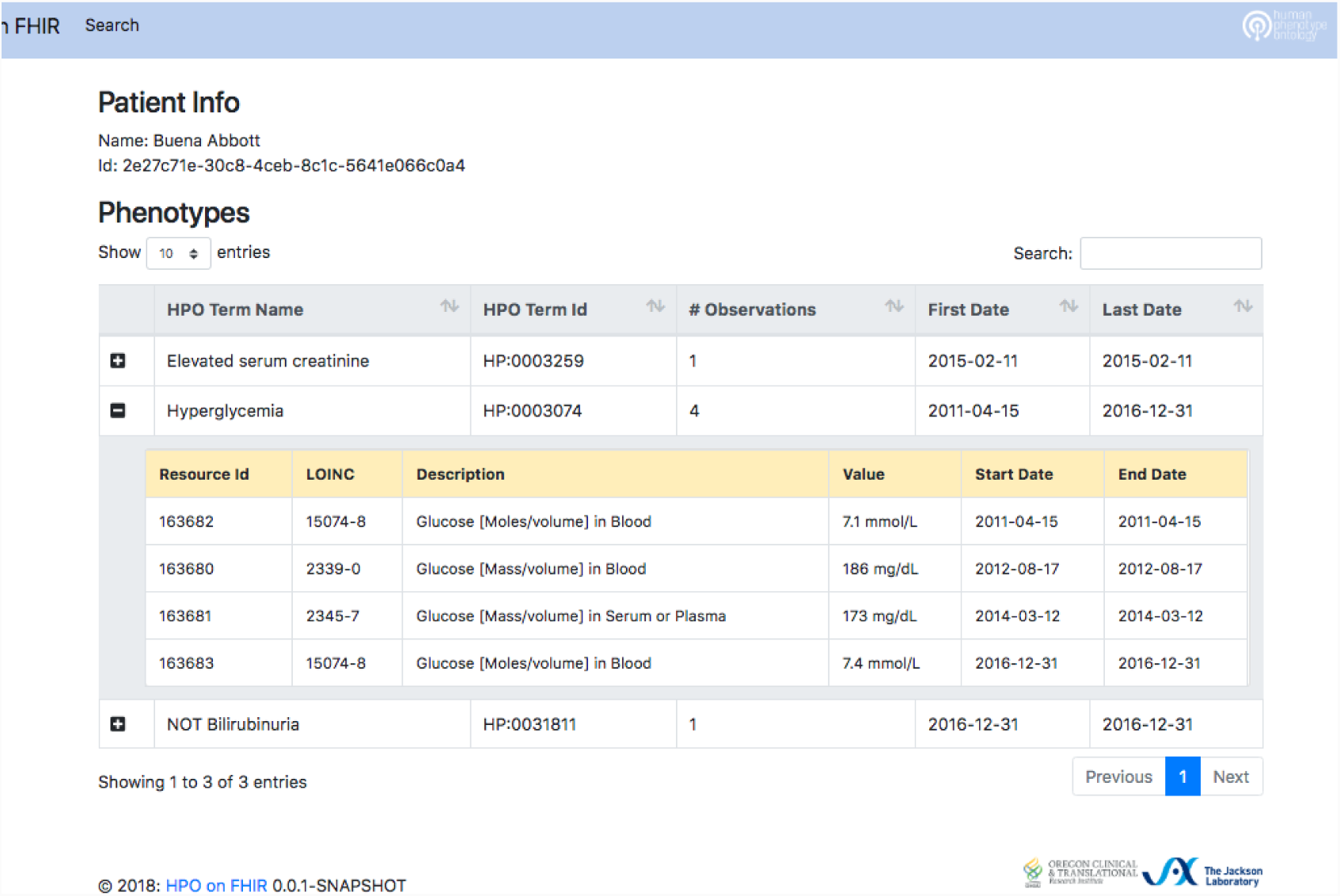
Screenshot of the HPO on FHIR app. We connected the HPO on FHIR app to the SmartHealthIT R3 Sandbox (a test server with synthetic data), queried all laboratory tests related to a simulated patient and converted all the laboratory tests into the corresponding HPO terms. The column “# Observations” shows the counts of laboratory tests that were mapped to the same HPO term, and for multiple tests mapped to the same HPO term, the dates of the first and last test are shown. In this example, LOINC 15074-8 was performed twice, and LOINC 2339-0 and 2345-7 were each performed once; the outcomes of all four tests were abnormally high (blood glucose), and so all four outcomes were mapped to the HPO term *Hyperglycemia* (HP:0003074).

### LOINC to HPO demonstration with asthma

To test our method for semantic integration of laboratory tests, we analyzed a de-identified EHR dataset from the University of North Carolina (UNC) comprised of 15,681 patients that had a history of asthma or asthma-like symptoms. The cohort is skewed toward female (58.9%) and older patients (median age: 61.5 years, Fig. 4A). The median tracking period of patients in this cohort is 3.1 years. The dataset contains ~54 million records of LOINC-coded clinical test results, medication prescriptions, diagnosis codes, procedure codes, patient information and other supporting records (Fig. 4B). Using our LOINC to HPO conversion algorithm, we successfully transformed 9.3 out of 11 million (83.1%) laboratory tests into HPO terms (Fig. 4C). For the entire cohort, on average, each HPO term was mapped from 1.8 distinct types of laboratory tests (Fig. 4D), indicating that the transformation successfully integrated distinctly coded laboratory tests that have the same clinical interpretation. The mapping procedure assigned an average of 594 laboratory test-derived HPO terms per individual patient, many of which were from the same laboratory tests performed at different visits. The tests corresponded to a mean of 53.5 unique HPO terms, of which 20.8 were abnormalities and the remainder were normal phenotypes (Fig. 4E). The hierarchical structure of the HPO allows inferences to be propagated up to parent terms and their ancestors^13^; using this method, we inferred an additional 46.1 HPO terms (total 66.9) based on 20.8 abnormalities to each patient (Fig. S2).

**Fig. 4.**
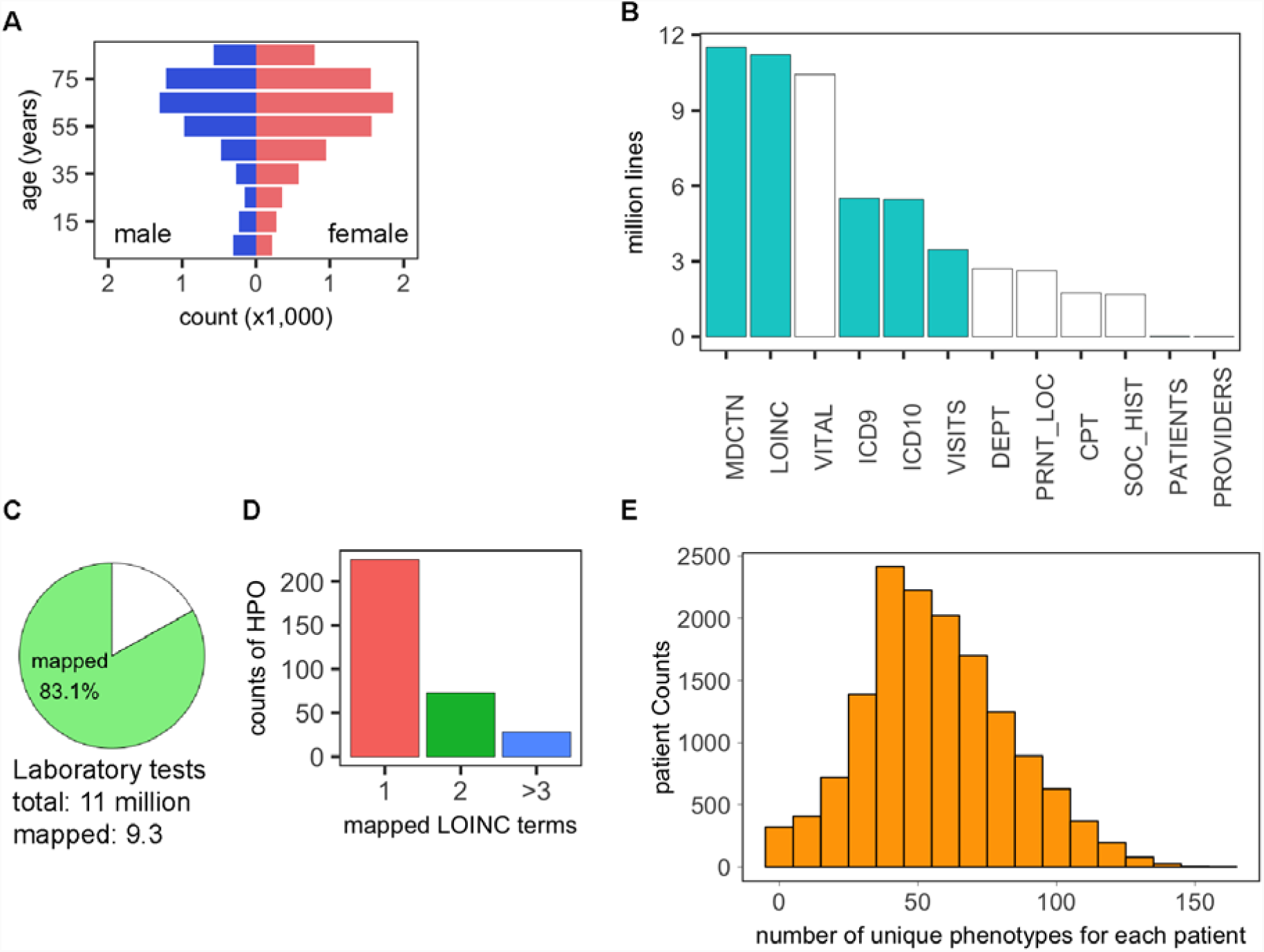
Analysis of UNC asthma dataset on asthma- and asthma-like patients. **A**. Age and sex distribution of patients. **B.** Categories of information extracted from the EHR data. Cyan, used for current research; white, not used for current research. MDCTN, medication; LOINC, LOINC-coded laboratory tests; VITAL, vital signs; ICD9, ICD 9-coded diagnosis; ICD10, ICD 10-coded diagnosis; VISITS, patient visit records; DEPT, clinic location; PRNT_LOC, hospital location; CPT, CPT-coded procedures; SOC_HIST, social history; PATIENTS, patient identification records; PROVIDER, provider information. **C.** Percentage of laboratory tests in our dataset that could be converted into HPO terms (the remaining unmapped tests did not have LOINC to HPO annotations). **D.** Number of LOINC terms mapped to a given HPO term in the UNC dataset. **E.** Distribution of patients by the number of unique HPO terms that are mapped to each patient.

As a proof-of-principle, we tested the ability of our procedure to identify phenotypic abnormalities associated with a diagnosis of asthma or with frequent prednisone use. About one third of the patients in this cohort had an ICD-9/10 diagnosis of asthma, and the remaining patients had ICD-9/10 codes reflecting other, potentially asthma-like, respiratory complaints. 14.2% of patients that had a diagnosis of asthma were administered or prescribed prednisone >3 times within a tracking period between 2004-2016; 8.5% of the remaining patients had been administered prednisone more than three times. Prednisone is a corticosteroid drug used for severe asthma treatment with multiple other indications^14^. We reasoned that both the diagnosis of asthma and the history of treatment with prednisone would likely be correlated with different but overlapping sets of laboratory abnormalities. Using logistic regression, we assessed the contribution of frequent prednisone prescription and the presence of acute asthma diagnosis to each phenotypic abnormality. Prednisone usage was significantly associated with an increased odds ratio for exhibiting many abnormal phenotypes that are consistent with the known effects of prednisone (Table 2), such as *Hypoalbuminemia* (HP:0003073)^15^, *Neutrophilia* (HP:0011897)^16^, *Monocytosis* (HP:0012311)^17^, *Leukocytosis* (HP:0001974)^17^, *Hypokalemia* (HP:0002900)^18^, and *Elevated serum creatine phosphokinase* (HP:0003236)^19^. An acute asthma diagnosis was significantly associated with five phenotypes, *Increased red blood cell count* (HP:0020059), *Increased VLDL cholesterol concentration* (HP:0003362) and *Eosinophilia* (HP:0001880), and two ancestor terms of *Eosinophilia, Abnormal eosinophil count* (HP:0020064) and *Abnormal eosinophil morphology* (HP:0001879). Eosinophilia is a well established marker for acute allergic asthma^20^. Although there have been some conflicting results^21^, a number of studies have shown a positive correlation between increased total, high or low-high-density lipoprotein cholesterol, or triglycerides and asthma^22–26^. An increased red blood cell count is not a recognized biomarker of asthma, but could conceivably reflect a number of factors including hypoxemia (11.1% with an acute asthma diagnosis also had a chronic obstructive pulmonary disease diagnosis), or hemoconcentration resulting from acute dehydration during an asthma attack, but the nature of this retrospective study does not allow us to consult the full medical records to investigate this.

**Table 2.**
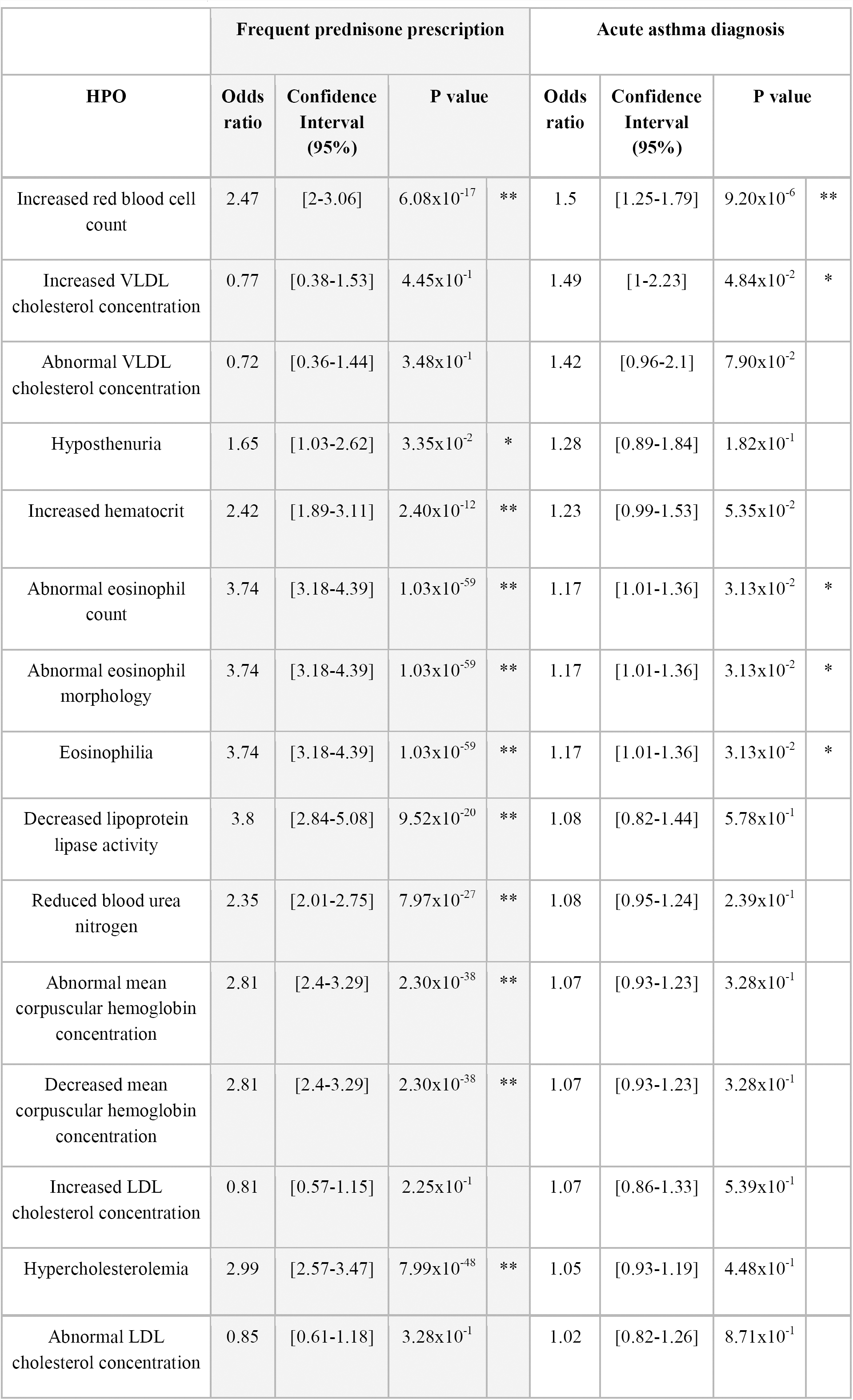
Odds ratio of phenotypes for frequent prednisone prescription and acute asthma diagnosis (*: p < 0.05, **: p < 0.01; table is sorted by the odds ratio for acute asthma diagnosis. Only HPO terms of which the odds ratio > 1 for acute asthma diagnosis are shown. Refer to Table S3 for all terms)

|  |  |  |  |  |  |  |  |  |
| --- | --- | --- | --- | --- | --- | --- | --- | --- |
| Increased red blood cell count | 2.47 | [2-3.06] | $6.08 \times 10^{-17}$ | ** | 1.5 | [1.25-1.79] | $9.20 \times 10^{-6}$ | ** |
| Increased VLDL cholesterol concentration | 0.77 | [0.38-1.53] | $4.45 \times 10^{-1}$ | | 1.49 | [1-2.23] | $4.84 \times 10^{-2}$ | * |
| Abnormal VLDL cholesterol concentration | 0.72 | [0.36-1.44] | $3.48 \times 10^{-1}$ | | 1.42 | [0.96-2.1] | $7.90 \times 10^{-2}$ | |
| Hyposthenuria | 1.65 | [1.03-2.62] | $3.35 \times 10^{-2}$ | * | 1.28 | [0.89-1.84] | $1.82 \times 10^{-1}$ | |
| Increased hematocrit | 2.42 | [1.89-3.11] | $2.40 \times 10^{-12}$ | ** | 1.23 | [0.99-1.53] | $5.35 \times 10^{-2}$ | |
| Abnormal eosinophil count | 3.74 | [3.18-4.39] | $1.03 \times 10^{-59}$ | ** | 1.17 | [1.01-1.36] | $3.13 \times 10^{-2}$ | * |
| Abnormal eosinophil morphology | 3.74 | [3.18-4.39] | $1.03 \times 10^{-59}$ | ** | 1.17 | [1.01-1.36] | $3.13 \times 10^{-2}$ | * |
| Eosinophilia | 3.74 | [3.18-4.39] | $1.03 \times 10^{-59}$ | ** | 1.17 | [1.01-1.36] | $3.13 \times 10^{-2}$ | * |
| Decreased lipoprotein lipase activity | 3.8 | [2.84-5.08] | $9.52 \times 10^{-20}$ | ** | 1.08 | [0.82-1.44] | $5.78 \times 10^{-1}$ | |
| Reduced blood urea nitrogen | 2.35 | [2.01-2.75] | $7.97 \times 10^{-27}$ | ** | 1.08 | [0.95-1.24] | $2.39 \times 10^{-1}$ | |
| Abnormal mean corpuscular hemoglobin concentration | 2.81 | [2.4-3.29] | $2.30 \times 10^{-38}$ | ** | 1.07 | [0.93-1.23] | $3.28 \times 10^{-1}$ | |
| Decreased mean corpuscular hemoglobin concentration | 2.81 | [2.4-3.29] | $2.30 \times 10^{-38}$ | ** | 1.07 | [0.93-1.23] | $3.28 \times 10^{-1}$ | |
| Increased LDL cholesterol concentration | 0.81 | [0.57-1.15] | $2.25 \times 10^{-1}$ | | 1.07 | [0.86-1.33] | $5.39 \times 10^{-1}$ | |
| Hypercholesterolemia | 2.99 | [2.57-3.47] | $7.99 \times 10^{-48}$ | ** | 1.05 | [0.93-1.19] | $4.48 \times 10^{-1}$ | |
| Abnormal LDL cholesterol concentration | 0.85 | [0.61-1.18] | $3.28 \times 10^{-1}$ | | 1.02 | [0.82-1.26] | $8.71 \times 10^{-1}$ | |

## Discussion

In this report, we present an approach to the semantic integration of laboratory tests and results in EHR data. Our approach connects a widely used system for denoting laboratory tests, LOINC, with a current standard for transmitting health care information, FHIR, and a computational resource for deep phenotyping that was previously used mainly in the context of rare disease research and diagnostics. Normalizing laboratory tests with HPO terms is an effective solution for two fundamental issues in clinical research: data integration and deep phenotyping. Laboratory test results support a large proportion of medical decisions^27^. It is commonplace that different laboratory tests may lead to results that have very similar or identical clinical interpretations. These different tests are recorded in the EHR using distinct codes (for instance, currently, there are four different LOINC terms for different tests of urine nitrite). This level of granularity can create difficulties for the semantic integration of comparable test results. By converting the results of laboratory tests to HPO-encoded phenotypes, our method provides an effective way for integrating laboratory tests that have the same clinical interpretation but different LOINC codes. Extracted patient phenotypes can be directly utilized for PheWAS studies, which is important because phenotyping patients is a major bottleneck for conducting PheWAS studies^28^. The Electronic Medical Records and Genomics (eMerge) network develops EHR-derived phenotyping algorithms by combining diagnosis codes, procedure codes, medication, narratives and subsets of laboratory tests and iteratively refine them to identify control and disease cohorts for genome- and phenome-wide association studies^1,3,28–30^. Our method complements existing phenotyping algorithms because it extracts additional phenotypic information by systematically interrogating the vast amount of data in laboratory tests.

The analysis of UNC EHR data demonstrated the potential of combining deep phenotypes from our tool with EHR data for biomarker discovery. Our current mapping library allowed us to convert the majority of the laboratory tests into HPO terms and assign an average of 53.5 unique phenotypes to each patient. The statistical analysis identified phenotypic abnormalities that are associated with frequent prescriptions of prednisone and/or acute asthma diagnosis. The cohort used for this analysis is biased toward senior and female patients and may not be reflective of asthma patient distributions, but the fact that our analysis identified numerous abnormalities that are associated with either prednisone use or asthma suggests that our approach can be useful for the investigation of EHR data for laboratory-based biomarkers of diseases and conditions. We have demonstrated the utility of our approach on the UNC dataset using a simple logistic regression approach as a proof of principle; we envision that our mapping approach could be used together with a variety of statistical and algorithmic analysis strategies to address a variety of topics in EHR-based translational research, and we have therefore coded our foundational approach in a way that can easily be integrated into other statistical analysis pipelines.

Several other use cases for our approach are conceivable. Rule-based algorithms could be applied to infer HPO terms from the primary phenotypic abnormalities. For instance, the combination of *Decreased hemoglobin concentration* (HP:0020062) and *Decreased mean corpuscular volume* (HP:0025066) implies *Microcytic anemia* (HP:0001935). The HPO is widely used in rare disease diagnostics, but one bottleneck is that in many settings, HPO terms need to be entered manually into the analysis software. A recent study used text-mining to extract detailed patient phenotypes through natural language processing of clinical narratives in EHR, and used the resulting lists of HPO terms for genomic diagnostics^11^. Our tool could supplement such tools by providing a computational representation of laboratory findings. In principle, our tool could be used to support other tasks related to EHR data, including decision support and cohort recruitment. In the future, we anticipate that semantic integration of a wider range of EHR data will become the norm to support data-driven translational research and precision medicine.

## Methods

### Mapping LOINC terms to HPO terms

We performed manual biocuration to construct a mapping library from each potential outcome of a LOINC test to the corresponding HPO term (Fig. 1A). The test outcome is represented using a subset of FHIR codes (Table 1, primary code), such as ‘lower than normal’, ‘normal’, or ‘higher than normal’. For quantitative tests that report a numeric measurement, we use FHIR interpretation code “L” and “H” to indicate lower or higher than normal, and “N” and “A” to indicate the result is normal or abnormal. For ordinal tests that have a binary outcome, i.e. present or absent of the test target, we use FHIR interpretation code “POS” to indicate present and “NEG” to indicate absent. Additionally, other interpretation codes defined by FHIR are first mapped to primary codes. For example, FHIR codes “LL” (critically low) and “<” (off scale low) are both mapped to “L” (Table 1).

The value for a map entry is an HPO term accompanied by a boolean value to indicate whether it should be negated. That is, while an abnormal test outcome is mapped to a particular HPO term, the normal outcome for that test is mapped to the negated form, since the HPO contains only terms for abnormal phenotypes. Fig. 1A shows three examples of mappings for Qn, Ord or Nom LOINC terms.

### LOINC to HPO mapping file

The LOINC to HPO mapping file contains records of mapping from LOINC test outcomes to the corresponding HPO terms. The annotation data is serialized as a tab-separated value (TSV) file. Each line records the LOINC code, test outcome, the mapped HPO term, and whether the mapped term should be negated. The annotation file is deposited at Github and can be accessed at https://w3id.org/loinc2hpo/annotations. An excerpt is shown in Supplemental Table 1.

### HPO on FHIR

We created a SMART on FHIR application, HPO on FHIR, to query a FHIR-enabled EHR servers and return patient laboratory results with LOINC codes and their corresponding HPO terms. The web interface of the application aggregates identical HPO terms together for visualization and also allows users to display source laboratory tests including subject, LOINC code, FHIR resource id, effective time and the corresponding HPO term. The app was written in the Java language with the Spring framework. The app uses the LOINC to HPO conversion algorithm described in fig. S1. The app is deposited at Github and can be accessed at (https://github.com/OCTRI/poc-hpo-on-fhir).

### Command-Line application for gathering FHIR server statistics

We created a command-line application that finds all laboratory tests for a patient on a FHIR server and attempts to convert them to HPO. The conversion results, both successes and failures, are stored in a relational database to aid in translational research. We ran the application on 7 common FHIR sandboxes and gathered statistics about the LOINCs encountered, the rate of success in conversion, and the underlying causes of failure. The application was written in the Java language with the Spring framework. Source code, results, and a backup of the database, can be accessed at (https://github.com/OCTRI/f2hstats).

### Analysis of UNC data on patients with asthma or an asthma-like condition

For the purposes of demonstrating the potential utility of our library, we examined a deidentified EHR dataset extracted from the Carolina Data Warehouse for Health (CDWH) at the University of North Carolina (UNC). The data was accessed under a fully executed Data Use Agreement between The Jackson Laboratory and UNC. The CDWH is UNC Health Care System’s (UNCHCS) enterprise data warehouse, and contains EHR data for all UNCHCS patients from 2004 through 2016. The sample used for this investigation contains 15681 patients with one or more encounters at UNCHCS with an asthma or asthma-like diagnosis (Table S2). The data was exported from the UNC EHR system as 8 separate comma-separated value (CSV) files containing clinical observations in a variety of data domains, including demographics, encounter details, diagnoses, procedures, medications, vital signs, and LOINC-coded lab results. Prior to transmission from UNC, the dataset was deidentified according to the Safe Harbor method of the Health Insurance Portability and Accountability Act (HIPAA), and all dates were shifted +/-50 days.

Using the extracted laboratory data, we converted each LOINC-coded test into an HPO term. We note, however, that not every laboratory test result was captured in the available dataset. For each patient, we combined test records mapped to the same HPO terms and recorded the counts of observations for each HPO term. Then we inferred additional phenotypic abnormalities based on the hierarchical structure of HPO, i.e. if a patient was assigned with an HPO term, we infer that the patient automatically had phenotypic abnormalities encoded by parent and other ancestor terms (fig. S2). We reasoned that an isolated abnormal measurement might represent an artefact or might not be typical of the clinical course of the patient, and therefore used a threshold of 3 observations over the entire observation period in order to classify a patient with the corresponding HPO-encoded phenotypic abnormality. We classified a patient not having an HPO-encoded phenotypic abnormality only when the patient had never been assigned for such an HPO term. Patient age was calculated from the last hospital visit date subtracting the birth date and is subject to an inaccuracy of +/- 50 days due to the deidentification procedure (see above). Patients that rarely visited hospitals were less likely to receive laboratory tests and thus had less phenotypes, so we excluded those that had medical encounters on less than 10 days. Patients received more than 3 prednisone prescriptions were considered frequent users.

## Statistics

We applied logistic regression model to determine the weights of being frequent prednisone user (values 0 or 1) and having acute asthma diagnosis (values 0 or 1) in determining a patient having an HPO-encoded phenotype (values 0 or 1). We excluded HPO terms from analysis of which the majority (95%) of the cohort had universal values (all 0 or 1). The natural exponential of the weights ± 1.98 standard deviations were converted to the odd ratio and 95% confidence intervals for each variable.

Data cleaning, normalization, wrangling and table joining were conducted by a combination of “tidyverse”, “RSQLite” packages in R, SQLite and Java. Logistic regression was conducted with the “glm” package in R. All source code is deposited at Github and can be accessed through https://github.com/TheJacksonLaboratory/HUSHDataAnalysis.

## List of Supplementary Materials

Fig. S1. FHIR to HPO conversion algorithm.

Fig. S2. Inference of phenotypic abnormalities with the hierarchy of HPO.

Table S1. Excerpt of LOINC to HPO annotation file.

Table S2. ICD codes used by UNC asthma dataset to identify asthma and asthma-like patients.

Table S3. Odds ratio of phenotypes for frequent prednisone prescription and acute asthma diagnosis

## Acknowledgments

The authors acknowledge colleagues from the Monarch Initiative for comments on this project. Research reported in this work was supported by the National Institutes of Health’s National Center for Advancing Translational Sciences, Grant Number U24TR00230, the Biomedical Data Translator program (awards OT3TR002019 and OT3TR002020), and the Clinical and Translational Science program (award UL1TR002489). The project also received support from the Intramural Research Program within the National Library of Medicine, National Institutes of Health and the National Human Genome Research Institute, National Institutes of Health (award NR24OD011883). This work was also supported by the U.S. National Library of Medicine contract HHSN276201400008C. The content is solely the responsibility of the authors and does not necessarily represent the official views of the National Institutes of Health. This material contains content from LOINC^®^ (http://loinc.org) which is copyright © 1995-2018, Regenstrief Institute, Inc. and the Logical Observation Identifiers Names and Codes (LOINC) Committee and is available at no cost under the license at http://loinc.org/license.

## Author contributions

XAZ: software engineering, curation, data analysis, and interpretation; AY: software engineering; NV, JPG, LCC, PNR: data curation; DD: software engineering; MPJ, VR, KR: data analysis; ERP,JC,KF: data collection and interpretation; DBP, HX, RZ, JR, NAW, SK, CM, DV, CGC, CMl, MAH: data interpretation; XAZ, PNR designed study, wrote manuscript.

## Data and materials availability

The patient EHR dataset can be acquired from the UNC with Data Use Agreement.

## References

1. Denny, J. C., Bastarache, L. & Roden, D. M. Phenome-Wide Association Studies as a Tool to Advance Precision Medicine. Annu. Rev. Genomics Hum. Genet. 17, 353–373 (2016).

2. Verma, A. et al. PheWAS and Beyond: The Landscape of Associations with Medical Diagnoses and Clinical Measures across 38,662 Individuals from Geisinger. Am. J. Hum. Genet. 102, 592–608 (2018).

3. Denny, J. C. et al. Variants near FOXE1 are associated with hypothyroidism and other thyroid conditions: using electronic medical records for genome- and phenome-wide studies. Am. J. Hum. Genet. 89, 529–542 (2011).

4. Dewey, F. E. et al. Distribution and clinical impact of functional variants in 50,726 whole-exome sequences from the DiscovEHR study. Science 354, aaf6814 (2016).

5. Freimer, N. & Sabatti, C. The human phenome project. Nat. Genet. 34, 15–21 (2003).

6. Robinson, P. N. Deep phenotyping for precision medicine. Hum. Mutat. 33, 777–780 (2012).

7. Leroux, H., Metke-Jimenez, A. & Lawley, M. J. Towards achieving semantic interoperability of clinical study data with FHIR. J. Biomed. Semantics 8, 41 (2017).

8. McDonald, C. J. et al. LOINC, a universal standard for identifying laboratory observations: a 5-year update. Clin. Chem. 49, 624–633 (2003).

9. Köhler, S. et al. The Human Phenotype Ontology in 2017. Nucleic Acids Res. 45, D865–D876 (2017).

10. Posey, J. E. et al. Resolution of Disease Phenotypes Resulting from Multilocus Genomic Variation. N. Engl. J. Med. 376, 21–31 (2017).

11. Son, J. H. et al. Deep Phenotyping on Electronic Health Records Facilitates Genetic Diagnosis by Clinical Exomes. Am. J. Hum. Genet. 103, 58–73 (2018).

12. Mandel, J. C., Kreda, D. A., Mandl, K. D., Kohane, I. S. & Ramoni, R. B. SMART on FHIR: a standards-based, interoperable apps platform for electronic health records. J. Am. Med. Inform. Assoc. 23, 899–908 (2016).

13. Robinson, P. N. & Bauer, S. Introduction to Bio-Ontologies. (CRC Press Inc, 2011).

14. Krishnan, J. A., Davis, S. Q., Naureckas, E. T., Gibson, P. & Rowe, B. H. An umbrella review: corticosteroid therapy for adults with acute asthma. Am. J. Med. 122, 977–991 (2009).

15. Aplasca, E. C. & Rammohan, M. The effect of prednisone on the levels of serum albumin of 20 patients with renal transplants. J. Am. Diet. Assoc. 86, 1404–1405 (1986).

16. Dale, D. C., Fauci, A. S., Guerry D I. V. & Wolff, S. M. Comparison of agents producing a neutrophilic leukocytosis in man. Hydrocortisone, prednisone, endotoxin, and etiocholanolone. J. Clin. Invest. 56, 808–813 (1975).

17. Shoenfeld, Y., Gurewich, Y., Gallant, L. A. & Pinkhas, J. Prednisone-induced leukocytosis. Influence of dosage, method and duration of administration on the degree of leukocytosis. Am. J. Med. 71, 773–778 (1981).

18. Veltri, K. T. & Mason, C. Medication-induced hypokalemia. P T 40, 185–190 (2015).

19. Smithson, J. & Others. Drug induced muscle disorders. Australian pharmacist 28, 1056 (2009).

20. Price, D. B. et al. Blood eosinophil count and prospective annual asthma disease burden: a UK cohort study. Lancet Respir Med 3, 849–858 (2015).

21. Yiallouros, P. K. et al. Low serum high-density lipoprotein cholesterol in childhood is associated with adolescent asthma. Clin. Exp. Allergy 42, 423–432 (2012).

22. Ramaraju, K., Krishnamurthy, S., Maamidi, S., Kaza, A. M. & Balasubramaniam, N. Is serum cholesterol a risk factor for asthma? Lung India 30, 295–301 (2013).

23. Ko, S.-H. et al. Lipid profiles in adolescents with and without asthma: Korea National Health and nutrition examination survey data. Lipids Health Dis. 17, 158 (2018).

24. Chen, Y. C. et al. Lipid profiles in children with and without asthma: interaction of asthma and obesity on hyperlipidemia. Diabetes Metab. Syndr. 7, 20–25 (2013).

25. Al-Shawwa, B., Al-Huniti, N., Titus, G. & Abu-Hasan, M. Hypercholesterolemia is a potential risk factor for asthma. J. Asthma 43, 231–233 (2006).

26. Cottrell, L., Neal, W. A., Ice, C., Perez, M. K. & Piedimonte, G. Metabolic abnormalities in children with asthma. Am. J. Respir. Crit. Care Med. 183, 441–448 (2011).

27. Badrick, T. Evidence-based laboratory medicine. Clin. Biochem. Rev. 34, 43–46 (2013).

28. Pathak, J., Kho, A. N. & Denny, J. C. Electronic health records-driven phenotyping: challenges, recent advances, and perspectives. J. Am. Med. Inform. Assoc. 20, e206–11 (2013).

29. Ritchie, M. D. et al. Genome- and phenome-wide analyses of cardiac conduction identifies markers of arrhythmia risk. Circulation 127, 1377–1385 (2013).

30. Karnes, J. H. et al. Phenome-wide scanning identifies multiple diseases and disease severity phenotypes associated with HLA variants. Sci. Transl. Med. 9, (2017).

