## Supplementary materials for "Semantic Integration of Clinical Laboratory Tests from Electronic Health Records for Deep Phenotyping and Biomarker Discovery"

**Fig. S1**. FHIR to HPO conversion algorithm. Briefly, we extract the LOINC codes in the FHIR message, interpret the results with a code that is used in the mapping library, and then return the corresponding HPO term. If the result of a FHIR message is provided as a code used in the mapping library (such as the color of urine), the interpretation step is skipped; if the result of a FHIR message is provided with an interpretation code, the algorithm converts the interpretation code into a FHIR code used in Table 1; if the result is provided as a text string, the algorithm parses it and converts it to a FHIR code. For other tests, the algorithm compares the raw result with the reference ranges and assigns a code to represent the outcome. The algorithm was implemented as a Java library, fhir2hpo, with the Spring framework and can be accessed through the Github repository at <https://github.com/OCTRI/fhir2hpo>.

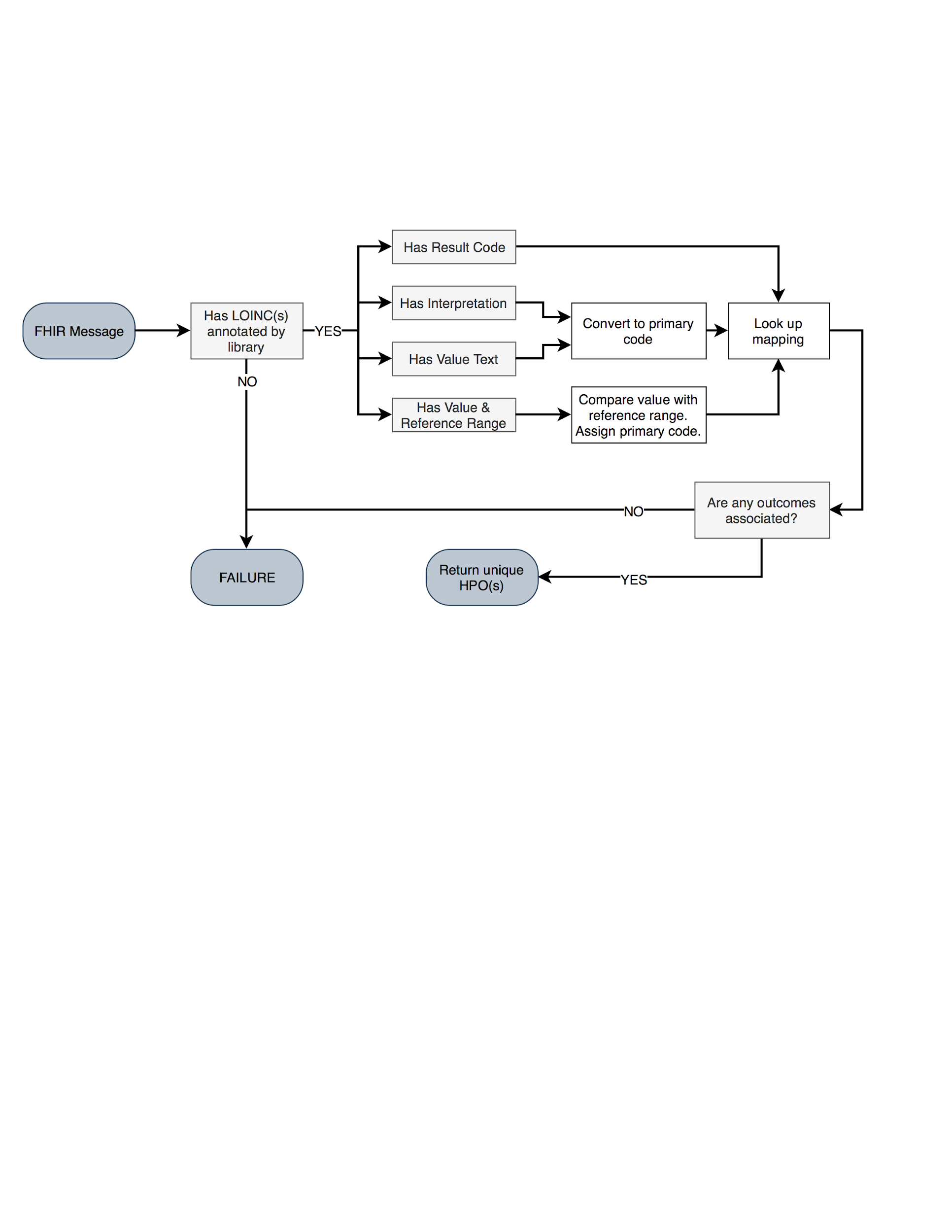

**Fig. S2.** Inference of phenotypic abnormalities with the hierarchy of HPO. The diagram demonstrates the tree-structured hierarchy with a few selected terms related to very low density lipoprotein (VLDL) cholesterol. The leaf terms contain more specific information than the parent term. We infer, for instance, that if a patient has the phenotype of *Increased VLDL cholesterol concentration* (HP:0003362), the patient also has abnormal phenotypes encoded by the ancestors of the current term, such as *Abnormal VLDL cholesterol concentration* (HP:0031889), *Abnormality of lipoprotein cholesterol concentration* (HP:0010979) and *Abnormality of cholesterol metabolism* (HP:0003107).

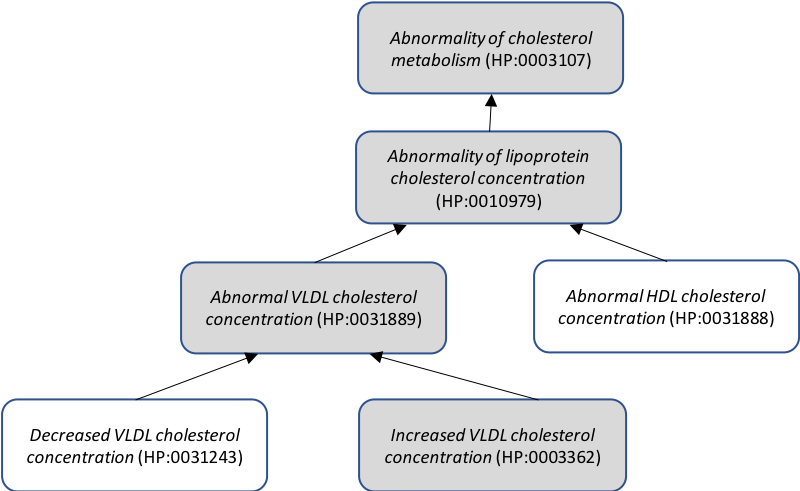

**Table S1**. Excerpt of LOINC to HPO annotation file. Columns from left to right: loincId, LOINC code; loincScale, type of the laboratory test; system and code, indicating the namespace and code for the test outcome; hpoTermId and isNegated, representing the mapped HPO and whether it needs to be negated for the outcome. Meta-data columns can be accessed online and are not shown here.

| **loincId** | **loincScale** | **system** | **code** | **hpoTermId** | **isNegated** |
| --- | --- | --- | --- | --- | --- |
| 38230-9 | Qn | FHIR | N | HP:0040077 | true |
| 38230-9 | Qn | FHIR | H | HP:0003072 | false |
| 38230-9 | Qn | FHIR | L | HP:0002901 | false |
| 5902-2 | Qn | FHIR | N | HP:0008151 | true |
| 5902-2 | Qn | FHIR | H | HP:0008151 | false |
| 777-3 | Qn | FHIR | N | HP:0011873 | true |
| 777-3 | Qn | FHIR | H | HP:0001894 | false |
| 777-3 | Qn | FHIR | L | HP:0001873 | false |
| 2019-8 | Qn | FHIR | H | HP:0012416 | false |
| 2019-8 | Qn | FHIR | L | HP:0012417 | false |
| 5803-2 | Qn | FHIR | L | HP:0031033 | false |
| ... | ... | ... | ... | ... | ... |

**Table S2**. ICD codes used by UNC asthma dataset to identify asthma and asthma-like patients. Patients were collected into this dataset if they have one of the ICD codes.

| **asthma diagnosis** | | **Asthma-like diagnosis** | |
| --- | --- | --- | --- |
| ICD-9: 493 | asthma | ICD-9:464 | croup |
| ICD-10: J45 | asthma | ICD-10:J05 | croup |
|  |  | ICD-9:496 | Reactive airway |
|  |  | ICD-10:J44 | Reactive airway |
|  |  | ICD-10:J66 | Reactive airway |
|  |  | ICD-9: 786 | cough |
|  |  | ICD-10: R05 | cough |
|  |  | ICD-9: 481-486 | pneumonia |
|  |  | ICD-10: J12-J18 | pneumonia |

**Table S3**. Odds ratio of phenotypes for frequent prednisone prescription and acute asthma diagnosis (asterisk: p < 0.05, double asterisk: p < 0·01; table is sorted by the odds ratio for acute asthma diagnosis)

|  | **Frequent prednisone prescription** | | | | **Acute asthma diagnosis** | | | |
| --- | --- | --- | --- | --- | --- | --- | --- | --- |
| **HPO** | **Odds**  **ratio** | **Confidence Interval (95%)** | **P value** | | **Odds**  **ratio** | **Confidence Interval (95%)** | **P value** | |
| Increased red blood cell count | 2·47 | [2-3·06] | 6·08x10^-17^ | ** | 1·5 | [1·25-1·79] | 9·20x10^-6^ | ** |
| Increased VLDL cholesterol concentration | 0·77 | [0·38-1·53] | 4·45x10^-1^ |  | 1·49 | [1-2·23] | 4·84x10^-2^ | * |
| Abnormal VLDL cholesterol concentration | 0·72 | [0·36-1·44] | 3·48x10^-1^ |  | 1·42 | [0·96-2·1] | 7·90x10^-2^ |  |
| Hyposthenuria | 1·65 | [1·03-2·62] | 3·35x10^-2^ | * | 1·28 | [0·89-1·84] | 1·82x10^-1^ |  |
| Increased hematocrit | 2·42 | [1·89-3·11] | 2·40x10^-12^ | ** | 1·23 | [0·99-1·53] | 5·35x10^-2^ |  |
| Abnormal eosinophil count | 3·74 | [3·18-4·39] | 1·03x10^-59^ | ** | 1·17 | [1·01-1·36] | 3·13x10^-2^ | * |
| Abnormal eosinophil morphology | 3·74 | [3·18-4·39] | 1·03x10^-59^ | ** | 1·17 | [1·01-1·36] | 3·13x10^-2^ | * |
| Eosinophilia | 3·74 | [3·18-4·39] | 1·03x10^-59^ | ** | 1·17 | [1·01-1·36] | 3·13x10^-2^ | * |
| Decreased lipoprotein lipase activity | 3·8 | [2·84-5·08] | 9·52x10^-20^ | ** | 1·08 | [0·82-1·44] | 5·78x10^-1^ |  |
| Reduced blood urea nitrogen | 2·35 | [2·01-2·75] | 7·97x10^-27^ | ** | 1·08 | [0·95-1·24] | 2·39x10^-1^ |  |
| Abnormal mean corpuscular hemoglobin concentration | 2·81 | [2·4-3·29] | 2·30x10^-38^ | ** | 1·07 | [0·93-1·23] | 3·28x10^-1^ |  |
| Decreased mean corpuscular hemoglobin concentration | 2·81 | [2·4-3·29] | 2·30x10^-38^ | ** | 1·07 | [0·93-1·23] | 3·28x10^-1^ |  |
| Increased LDL cholesterol concentration | 0·81 | [0·57-1·15] | 2·25x10^-1^ |  | 1·07 | [0·86-1·33] | 5·39x10^-1^ |  |
| Hypercholesterolemia | 2·99 | [2·57-3·47] | 7·99x10^-48^ | ** | 1·05 | [0·93-1·19] | 4·48x10^-1^ |  |
| Abnormal LDL cholesterol concentration | 0·85 | [0·61-1·18] | 3·28x10^-1^ |  | 1·02 | [0·82-1·26] | 8·71x10^-1^ |  |
| Decreased mean corpuscular volume | 2·54 | [2·17-2·96] | 1·08x10^-32^ | ** | 0·99 | [0·86-1·13] | 8·47x10^-1^ |  |
| Elevated serum bicarbonate concentration | 2·75 | [2·25-3·36] | 1·85x10^-23^ | ** | 0·99 | [0·82-1·18] | 8·78x10^-1^ |  |
| Leukocytosis | 4·53 | [3·95-5·19] | 3·57x10^-107^ | ** | 0·98 | [0·89-1·07] | 6·09x10^-1^ |  |
| Thrombocytosis | 2·56 | [2·18-2·99] | 7·80x10^-32^ | ** | 0·98 | [0·85-1·12] | 7·28x10^-1^ |  |
| Decreased mean platelet volume | 4·53 | [3·98-5·15] | 2·84x10^-118^ | ** | 0·97 | [0·87-1·07] | 5·06x10^-1^ |  |
| Abnormal granulocyte morphology | 5·75 | [4·95-6·7] | 1·18x10^-115^ | ** | 0·94 | [0·85-1·03] | 1·66x10^-1^ |  |
| Abnormality of myeloid leukocytes | 5·75 | [4·95-6·7] | 1·18x10^-115^ | ** | 0·94 | [0·85-1·03] | 1·66x10^-1^ |  |
| Hypoxemia | 2·85 | [2·35-3·46] | 7·38x10^-27^ | ** | 0·94 | [0·79-1·12] | 4·92x10^-1^ |  |
| Abnormal hemoglobin | 2·05 | [1·79-2·35] | 4·46x10^-25^ | ** | 0·93 | [0·83-1·03] | 1·65x10^-1^ |  |
| Elevated hemoglobin A1c | 2·05 | [1·79-2·35] | 4·46x10^-25^ | ** | 0·93 | [0·83-1·03] | 1·65x10^-1^ |  |
| Abnormal erythrocyte sedimentation rate | 9·13 | [7·51-11·11] | 9·49x10^-111^ | ** | 0·92 | [0·75-1·14] | 4·54x10^-1^ |  |
| Abnormal liver morphology | 2·83 | [2·5-3·21] | 5·57x10^-60^ | ** | 0·92 | [0·83-1·01] | 8·79x10^-02^ |  |
| Abnormality of the abdominal organs | 2·83 | [2·5-3·21] | 5·57x10^-60^ | ** | 0·92 | [0·83-1·01] | 8·79x10^-2^ |  |
| Abnormality of the digestive system | 2·83 | [2·5-3·21] | 5·57x10^-60^ | ** | 0·92 | [0·83-1·01] | 8·79x10^-2^ |  |
| Abnormality of the liver | 2·83 | [2·5-3·21] | 5·57x10^-60^ | ** | 0·92 | [0·83-1·01] | 8·79x10^-2^ |  |
| Elevated erythrocyte sedimentation rate | 9·13 | [7·51-11·11] | 9·49x10^-111^ | ** | 0·92 | [0·75-1·14] | 4·54x10^-1^ |  |
| Elevated hepatic transaminase | 2·83 | [2·5-3·21] | 5·57x10^-60^ | ** | 0·92 | [0·83-1·01] | 8·79x10^-2^ |  |
| Elevated serum alanine aminotransferase | 2·89 | [2·51-3·31] | 7·70x10^-52^ | ** | 0·92 | [0·82-1·04] | 1·88x10^-1^ |  |
| Hyperlipidemia | 2·43 | [2·08-2·86] | 2·83x10^-28^ | ** | 0·92 | [0·81-1·05] | 2·24x10^-1^ |  |
| Hypertriglyceridemia | 2·43 | [2·08-2·86] | 2·83x10^-28^ | ** | 0·92 | [0·81-1·05] | 2·24x10^-1^ |  |
| Abnormal circulating thyroxine level | 3·76 | [2·97-4·75] | 5·95x10^-29^ | ** | 0·91 | [0·72-1·14] | 3·94x10^-1^ |  |
| Abnormal thyroid hormone level | 3·76 | [2·97-4·75] | 5·95x10^-29^ | ** | 0·91 | [0·72-1·14] | 3·94x10^-1^ |  |
| Abnormality of the thyroid gland | 3·76 | [2·97-4·75] | 5·95x10^-29^ | ** | 0·91 | [0·72-1·14] | 3·94x10^-1^ |  |
| Abnormality of thyroid physiology | 3·76 | [2·97-4·75] | 5·95x10^-29^ | ** | 0·91 | [0·72-1·14] | 3·94x10^-1^ |  |
| Abnormal platelet volume | 6·13 | [5·31-7·08] | 3·96x10^-138^ | ** | 0·9 | [0·82-0·99] | 2·68x10^-2^ | * |
| Abnormality of the immune system | 5·37 | [4·37-6·6] | 1·33x10^-58^ | ** | 0·9 | [0·82-0·99] | 2·75x10^-2^ | * |
| Elevated C-reactive protein level | 7·83 | [6·55-9·36] | 8·30x10^-115^ | ** | 0·9 | [0·74-1·09] | 2·65x10^-1^ |  |
| Hematuria | 4·68 | [3·88-5·64] | 2·89x10^-60^ | ** | 0·9 | [0·75-1·09] | 2·78x10^-1^ |  |
| Neutrophilia | 4·79 | [4·18-5·48] | 5·19x10^-116^ | ** | 0·89 | [0·81-0·99] | 2·42x10^-2^ | * |
| Abnormal leukocyte count | 5·2 | [4·26-6·35] | 4·65x10^-60^ | ** | 0·88 | [0·8-0·97] | 7·42x10^-3^ | ** |
| Abnormal leukocyte morphology | 5·11 | [4·19-6·24] | 7·15x10^-59^ | ** | 0·88 | [0·8-0·97] | 6·52x10^-3^ | ** |
| Abnormality of cellular immune system | 5·11 | [4·19-6·24] | 7·15x10^-59^ | ** | 0·88 | [0·8-0·97] | 6·52x10^-3^ | ** |
| Abnormality of urine homeostasis | 3·42 | [3-3·9] | 4·83x10^-77^ | ** | 0·88 | [0·8-0·97] | 8·92x10^-3^ | ** |
| Hypokalemia | 2·81 | [2·47-3·2] | 1·31x10^-55^ | ** | 0·88 | [0·79-0·98] | 1·41x10^-2^ | * |
| Decreased serum iron | 3·84 | [3·17-4·65] | 5·65x10^-44^ | ** | 0·87 | [0·72-1·05] | 1·43x10^-1^ |  |
| Abnormal neutrophil count | 5·64 | [4·88-6·52] | 5·62x10^-124^ | ** | 0·86 | [0·79-0·95] | 2·30x10^-3^ | ** |
| Abnormality of neutrophils | 5·64 | [4·88-6·52] | 5·62x10^-124^ | ** | 0·86 | [0·79-0·95] | 2·30x10^-3^ | ** |
| Decreased serum creatinine | 1·69 | [1·48-1·94] | 1·65x10^-14^ | ** | 0·86 | [0·78-0·96] | 6·40x10^-3^ | ** |
| Hyperlipoproteinemia | 1·84 | [1·58-2·14] | 4·22x10^-15^ | ** | 0·86 | [0·76-0·97] | 1·12x10^-2^ | * |
| Increased circulating thyroxine level | 3·77 | [2·89-4·93] | 8·31x10^-23^ | ** | 0·86 | [0·65-1·12] | 2·58x10^-1^ |  |
| Abnormal urine cytology | 3·44 | [3·02-3·92] | 3·68x10^-78^ | ** | 0·85 | [0·77-0·94] | 2·14x10^-3^ | ** |
| Abnormality of blood and blood-forming tissues | 4·01 | [2·8-5·76] | 2·49x10^-14^ | ** | 0·85 | [0·73-0·98] | 2·84x10^-2^ | * |
| Monocytosis | 2·04 | [1·75-2·38] | 1·64x10^-20^ | ** | 0·85 | [0·75-0·96] | 1·08x10^-2^ | * |
| Abnormal serum iron | 4·24 | [3·55-5·07] | 7·67x10^-58^ | ** | 0·84 | [0·71-1·01] | 5·65x10^-2^ |  |
| Abnormality of the endocrine system | 3·52 | [2·88-4·3] | 2·24x10^-35^ | ** | 0·84 | [0·69-1·01] | 6·72x10^-2^ |  |
| Elevated serum anion gap | 2·8 | [2·34-3·36] | 1·36x10^-29^ | ** | 0·84 | [0·71-0·99] | 3·59x10^-2^ | * |
| Elevated serum aspartate aminotransferase | 2·99 | [2·62-3·43] | 3·17x10^-58^ | ** | 0·84 | [0·75-0·94] | 1·79x10^-3^ | ** |
| Pyuria | 3·47 | [3-4·03] | 3·06x10^-62^ | ** | 0·84 | [0·74-0·96] | 9·67x10^-3^ | ** |
| Abnormal enzyme/coenzyme activity | 3·43 | [3·02-3·9] | 3·78x10^-81^ | ** | 0·83 | [0·75-0·91] | 1·74x10^-4^ | ** |
| Hypercapnia | 2·81 | [2·46-3·21] | 1·72x10^-53^ | ** | 0·83 | [0·74-0·92] | 4·93x10^-4^ | ** |
| Abnormal erythrocyte morphology | 2·91 | [2·37-3·58] | 1·38x10^-24^ | ** | 0·82 | [0·74-0·91] | 1·68x10^-4^ | ** |
| Abnormality of alkaline phosphatase activity | 2·41 | [2·1-2·76] | 1·40x10^-37^ | ** | 0·82 | [0·74-0·92] | 6·25x10^-4^ | ** |
| Abnormality of circulating hormone level | 3·62 | [2·95-4·45] | 1·14x10^-35^ | ** | 0·82 | [0·67-1·01] | 5·48x10^-2^ |  |
| Abnormality of the genitourinary system | 3·91 | [3·41-4·5] | 6·72x10^-85^ | ** | 0·82 | [0·75-0·9] | 9·79x10^-6^ | ** |
| Abnormality of the urinary system | 3·91 | [3·41-4·5] | 6·72x10^-85^ | ** | 0·82 | [0·75-0·9] | 9·79x10^-6^ | ** |
| Abnormality of the urinary system physiology | 3·91 | [3·41-4·5] | 6·72x10^-85^ | ** | 0·82 | [0·75-0·9] | 9·79x10^-6^ | ** |
| Abnormality of immune system physiology | 6·66 | [5·39-8·23] | 1·61x10^-70^ | ** | 0·8 | [0·63-1] | 4·93x10^-2^ | * |
| Abnormality of potassium homeostasis | 3·59 | [3·15-4·1] | 5·90x10^-83^ | ** | 0·8 | [0·73-0·88] | 4·07x10^-6^ | ** |
| Abnormality of acid-base homeostasis | 3·65 | [3·2-4·16] | 8·00x10^-86^ | ** | 0·79 | [0·72-0·86] | 4·10x10^-7^ | ** |
| Abnormality of cholesterol metabolism | 2·47 | [2·17-2·82] | 2·89x10^-42^ | ** | 0·79 | [0·72-0·86] | 1·95x10^-7^ | ** |
| Hypernatremia | 3·22 | [2·69-3·85] | 1·57x10^-38^ | ** | 0·79 | [0·67-0·93] | 4·18x10^-3^ | ** |
| Abnormal blood carbon dioxide level | 3·97 | [3·47-4·53] | 2·27x10^-92^ | ** | 0·78 | [0·71-0·86] | 1·71x10^-7^ | ** |
| Abnormal levels of creatine kinase in blood | 3·04 | [2·56-3·61] | 1·04x10^-37^ | ** | 0·78 | [0·66-0·92] | 2·37x10^-3^ | ** |
| Abnormal mean corpuscular volume | 2·33 | [2·06-2·64] | 1·86x10^-41^ | ** | 0·78 | [0·7-0·86] | 7·11x10^-7^ | ** |
| Abnormal renal physiology | 4·03 | [3·55-4·58] | 9·33x10^-104^ | ** | 0·78 | [0·7-0·85] | 1·55x10^-7^ | ** |
| Abnormality of circulating enzyme level | 3·04 | [2·56-3·61] | 1·04x10^-37^ | ** | 0·78 | [0·66-0·92] | 2·37x10^-3^ | ** |
| Abnormality of the kidney | 4·03 | [3·55-4·58] | 9·33x10^-104^ | ** | 0·78 | [0·7-0·85] | 1·55x10^-7^ | ** |
| Abnormality of the upper urinary tract | 4·03 | [3·55-4·58] | 9·33x10^-104^ | ** | 0·78 | [0·7-0·85] | 1·55x10^-7^ | ** |
| Acidosis | 3·18 | [2·67-3·79] | 1·22x10^-39^ | ** | 0·78 | [0·66-0·92] | 3·43x10^-3^ | ** |
| Elevated alkaline phosphatase | 2·4 | [2·09-2·77] | 1·24x10^-34^ | ** | 0·78 | [0·69-0·88] | 6·43x10^-5^ | ** |
| Elevated serum creatine phosphokinase | 3·04 | [2·56-3·61] | 1·04x10^-37^ | ** | 0·78 | [0·66-0·92] | 2·37x10^-3^ | ** |
| Hyperglycemia | 1·04 | [0·92-1·18] | 5·40x10^-1^ |  | 0·78 | [0·72-0·85] | 2·35x10^-8^ | ** |
| Increased HDL cholesterol concentration | 2·64 | [2·22-3·14] | 6·17x10^-29^ | ** | 0·78 | [0·67-0·91] | 1·82x10^-3^ | ** |
| Increased serum lactate | 3·18 | [2·67-3·79] | 1·22x10^-39^ | ** | 0·78 | [0·66-0·92] | 3·43x10^-3^ | ** |
| Abnormal blood gas level | 3·87 | [3·38-4·42] | 9·70x10^-89^ | ** | 0·77 | [0·7-0·84] | 1·24x10^-8^ | ** |
| Abnormal glucose homeostasis | 1·12 | [0·99-1·27] | 8·07x10^-2^ |  | 0·77 | [0·7-0·84] | 1·78x10^-9^ | ** |
| Abnormal monocyte count | 2·96 | [2·58-3·38] | 9·96x10^-57^ | ** | 0·77 | [0·69-0·87] | 9·12x10^-6^ | ** |
| Abnormality monocyte morphology | 2·96 | [2·58-3·38] | 9·96x10^-57^ | ** | 0·77 | [0·69-0·87] | 9·12x10^-6^ | ** |
| Abnormality of blood glucose concentration | 1·11 | [0·98-1·26] | 9·08x10^-2^ |  | 0·77 | [0·7-0·84] | 1·86x10^-9^ | ** |
| Abnormality of carbohydrate metabolism/homeostasis | 1·12 | [0·99-1·27] | 8·07x10^-2^ |  | 0·77 | [0·7-0·84] | 1·78x10^-9^ | ** |
| Abnormality of iron homeostasis | 3·81 | [3·27-4·44] | 5·66x10^-68^ | ** | 0·77 | [0·66-0·88] | 2·08x10^-4^ | ** |
| Abnormality of lipid metabolism | 2·42 | [2·12-2·76] | 1·35x10^-39^ | ** | 0·77 | [0·71-0·84] | 2·45x10^-9^ | ** |
| Abnormality of the respiratory system | 3·87 | [3·38-4·42] | 9·70x10^-89^ | ** | 0·77 | [0·7-0·84] | 1·24x10^-8^ | ** |
| Functional respiratory abnormality | 3·87 | [3·38-4·42] | 9·70x10^-89^ | ** | 0·77 | [0·7-0·84] | 1·24x10^-8^ | ** |
| Hyperproteinemia | 2·81 | [2·28-3·47] | 2·61x10^-22^ | ** | 0·77 | [0·63-0·94] | 1·02x10^-2^ | * |
| Hypochloremia | 2·56 | [2·25-2·91] | 4·10x10^-47^ | ** | 0·77 | [0·69-0·85] | 7·15x10^-7^ | ** |
| Abnormal thrombocyte morphology | 4·56 | [3·9-5·32] | 1·42x10^-83^ | ** | 0·76 | [0·69-0·83] | 7·04x10^-10^ | ** |
| Abnormality of cation homeostasis | 3·44 | [2·91-4·06] | 1·13x10^-48^ | ** | 0·76 | [0·69-0·83] | 1·11x10^-9^ | ** |
| Abnormality of ion homeostasis | 3·83 | [3·15-4·66] | 4·95x10^-42^ | ** | 0·76 | [0·69-0·84] | 3·32x10^-8^ | ** |
| Abnormality of monovalent inorganic cation homeostasis | 3·59 | [3·12-4·14] | 6·80x10^-71^ | ** | 0·76 | [0·7-0·83] | 2·07x10^-9^ | ** |
| Abnormality of transition element cation homeostasis | 3·85 | [3·31-4·48] | 6·52x10^-70^ | ** | 0·76 | [0·66-0·87] | 1·04x10^-4^ | ** |
| Increased RBC distribution width | 3·09 | [2·73-3·49] | 2·27x10^-72^ | ** | 0·76 | [0·7-0·83] | 7·45x10^-10^ | ** |
| Abnormal serum anion gap | 3·38 | [2·96-3·85] | 5·11x10^-76^ | ** | 0·75 | [0·68-0·83] | 1·72x10^-8^ | ** |
| Abnormality of lipoprotein cholesterol concentration | 2·29 | [2·01-2·63] | 2·01x10^-34^ | ** | 0·75 | [0·68-0·83] | 1·24x10^-8^ | ** |
| Hypercalcemia | 2·79 | [2·25-3·46] | 3·20x10^-21^ | ** | 0·75 | [0·61-0·92] | 5·66x10^-3^ | ** |
| Hyperphosphatemia | 6·33 | [5·54-7·23] | 2·90x10^-166^ | ** | 0·75 | [0·67-0·85] | 5·08x10^-6^ | ** |
| Abnormal blood oxygen level | 2·59 | [2·22-3·02] | 1·71x10^-34^ | ** | 0·74 | [0·64-0·85] | 1·22x10^-5^ | ** |
| Abnormal hemoglobin concentration | 3·13 | [2·72-3·61] | 1·58x10^-56^ | ** | 0·74 | [0·68-0·81] | 5·69x10^-12^ | ** |
| Hypocapnia | 3·48 | [3·06-3·97] | 4·79x10^-80^ | ** | 0·74 | [0·67-0·83] | 1·17x10^-7^ | ** |
| Abnormal glomerular filtration rate | 3·71 | [3·28-4·19] | 3·82x10^-99^ | ** | 0·73 | [0·66-0·8] | 1·35x10^-10^ | ** |
| Abnormal hematocrit | 3·98 | [3·4-4·66] | 4·20x10^-67^ | ** | 0·73 | [0·66-0·79] | 6·79x10^-13^ | ** |
| Abnormal serum bicarbonate concentration | 3 | [2·57-3·5] | 1·23x10^-45^ | ** | 0·73 | [0·64-0·85] | 1·40x10^-5^ | ** |
| Abnormality of phosphate homeostasis | 5·74 | [5·04-6·54] | 2·93x10^-155^ | ** | 0·73 | [0·65-0·82] | 3·78x10^-8^ | ** |
| Decreased HDL cholesterol concentration | 1·98 | [1·66-2·35] | 6·00x10^-15^ | ** | 0·73 | [0·63-0·85] | 3·56x10^-5^ | ** |
| Decreased hemoglobin concentration | 2·98 | [2·6-3·41] | 2·84x10^-56^ | ** | 0·73 | [0·67-0·79] | 1·44x10^-13^ | ** |
| Hypolipoproteinemia | 1·98 | [1·67-2·34] | 8·49x10^-16^ | ** | 0·73 | [0·63-0·84] | 8·90x10^-6^ | ** |
| Abnormal circulating creatinine level | 2·71 | [2·37-3·1] | 2·52x10^-49^ | ** | 0·72 | [0·66-0·78] | 6·71x10^-14^ | ** |
| Abnormality of chloride homeostasis | 3·85 | [3·34-4·43] | 4·99x10^-80^ | ** | 0·72 | [0·66-0·79] | 1·82x10^-12^ | ** |
| Abnormality of divalent inorganic cation homeostasis | 3·18 | [2·78-3·64] | 1·76x10^-64^ | ** | 0·72 | [0·66-0·78] | 1·12x10^-13^ | ** |
| Decreased glomerular filtration rate | 3·71 | [3·28-4·2] | 2·77x10^-99^ | ** | 0·72 | [0·66-0·8] | 1·03x10^-10^ | ** |
| Elevated gamma-glutamyltransferase activity | 6·38 | [5·41-7·51] | 4·90x10^-111^ | ** | 0·72 | [0·61-0·86] | 2·83x10^-4^ | ** |
| Elevated serum creatinine | 2·49 | [2·21-2·82] | 3·62x10^-50^ | ** | 0·72 | [0·66-0·79] | 1·95x10^-12^ | ** |
| Hyperchloremia | 3·65 | [3·2-4·15] | 1·18x10^-87^ | ** | 0·72 | [0·65-0·79] | 6·39x10^-11^ | ** |
| Hypomagnesemia | 3·89 | [3·41-4·45] | 1·32x10^-89^ | ** | 0·72 | [0·64-0·82] | 1·94x10^-7^ | ** |
| Abnormality of magnesium homeostasis | 4·03 | [3·55-4·58] | 1·45x10^-104^ | ** | 0·71 | [0·64-0·79] | 6·98x10^-11^ | ** |
| Abnormality of nitrogen compound homeostasis | 2·98 | [2·54-3·5] | 1·42x10^-41^ | ** | 0·71 | [0·65-0·77] | 2·51x10^-14^ | ** |
| Abnormality of sodium homeostasis | 2·92 | [2·56-3·33] | 1·28x10^-58^ | ** | 0·71 | [0·64-0·78] | 2·25x10^-12^ | ** |
| Hyponatremia | 2·19 | [1·92-2·5] | 7·32x10^-33^ | ** | 0·71 | [0·64-0·79] | 6·13x10^-11^ | ** |
| Abnormal concentration of calcium in blood | 2·74 | [2·41-3·11] | 1·93x10^-54^ | ** | 0·7 | [0·64-0·76] | 1·40x10^-15^ | ** |
| Abnormal HDL cholesterol concentration | 2·91 | [2·53-3·35] | 5·03x10^-51^ | ** | 0·7 | [0·63-0·79] | 8·12x10^-10^ | ** |
| Abnormal red blood cell count | 3·66 | [3·15-4·25] | 8·81x10^-66^ | ** | 0·7 | [0·64-0·77] | 1·19x10^-15^ | ** |
| Abnormality of calcium homeostasis | 2·75 | [2·41-3·12] | 1·74x10^-54^ | ** | 0·7 | [0·64-0·76] | 2·06x10^-15^ | ** |
| Abnormality of circulating protein level | 5·62 | [4·87-6·48] | 2·03x10^-125^ | ** | 0·7 | [0·64-0·77] | 7·53x10^-14^ | ** |
| Hypermagnesemia | 4·03 | [3·5-4·64] | 5·09x10^-85^ | ** | 0·7 | [0·61-0·8] | 7·62x10^-8^ | ** |
| Hyperoxemia | 2·4 | [1·99-2·89] | 7·97x10^-21^ | ** | 0·7 | [0·59-0·83] | 2·78x10^-5^ | ** |
| Increased serum ferritin | 4·36 | [3·29-5·77] | 2·04x10^-25^ | ** | 0·7 | [0·52-0·95] | 1·91x10^-2^ | * |
| Reduced hematocrit | 3·75 | [3·24-4·34] | 1·67x10^-72^ | ** | 0·7 | [0·64-0·76] | 4·62x10^-17^ | ** |
| Decreased serum anion gap | 2·9 | [2·55-3·31] | 7·65x10^-58^ | ** | 0·69 | [0·62-0·77] | 9·34x10^-12^ | ** |
| Hypocalcemia | 2·54 | [2·24-2·87] | 8·22x10^-50^ | ** | 0·69 | [0·63-0·76] | 4·77x10^-16^ | ** |
| Hypoproteinemia | 3·48 | [3·05-3·96] | 6·11x10^-79^ | ** | 0·69 | [0·62-0·77] | 1·32x10^-11^ | ** |
| Abnormal lymphocyte morphology | 5·52 | [4·78-6·37] | 2·28x10^-123^ | ** | 0·68 | [0·61-0·74] | 3·51x10^-16^ | ** |
| Abnormal serum ferritin | 4·01 | [3·1-5·17] | 4·33x10^-27^ | ** | 0·68 | [0·52-0·9] | 5·18x10^-3^ | ** |
| Abnormal blood urea nitrogen | 2·49 | [2·19-2·84] | 5·15x10^-44^ | ** | 0·67 | [0·62-0·74] | 1·14x10^-18^ | ** |
| Azotemia | 2·42 | [2·13-2·76] | 5·85x10^-42^ | ** | 0·66 | [0·6-0·72] | 7·32x10^-22^ | ** |
| Hyperkalemia | 4·24 | [3·7-4·87] | 4·13x10^-96^ | ** | 0·66 | [0·58-0·75] | 8·07x10^-11^ | ** |
| Increased mean platelet volume | 3·5 | [3·01-4·07] | 6·11x10^-61^ | ** | 0·66 | [0·57-0·76] | 8·75x10^-9^ | ** |
| Increased mean corpuscular volume | 1·82 | [1·57-2·11] | 2·03x10^-15^ | ** | 0·65 | [0·57-0·74] | 3·75x10^-11^ | ** |
| Leukopenia | 5·2 | [4·51-5·99] | 6·92x10^-117^ | ** | 0·65 | [0·6-0·72] | 1·47x10^-19^ | ** |
| Abnormal lactate dehydrogenase activity | 4·74 | [3·92-5·75] | 2·64x10^-58^ | ** | 0·64 | [0·52-0·78] | 1·62x10^-5^ | ** |
| Conjugated hyperbilirubinemia | 2·53 | [1·99-3·22] | 1·78x10^-14^ | ** | 0·64 | [0·5-0·81] | 1·70x10^-4^ | ** |
| Hypoglycemia | 3·59 | [2·96-4·35] | 9·06x10^-40^ | ** | 0·64 | [0·52-0·77] | 5·74x10^-6^ | ** |
| Increased lactate dehydrogenase activity | 4·74 | [3·92-5·75] | 2·64x10^-58^ | ** | 0·64 | [0·52-0·78] | 1·62x10^-5^ | ** |
| Lymphopenia | 5·43 | [4·73-6·24] | 1·60x10^-128^ | ** | 0·64 | [0·58-0·7] | 3·42x10^-20^ | ** |
| Neutropenia | 3·24 | [2·79-3·76] | 4·48x10^-54^ | ** | 0·64 | [0·55-0·74] | 8·95x10^-10^ | ** |
| Decreased red blood cell count | 3·33 | [2·91-3·82] | 1·52x10^-68^ | ** | 0·63 | [0·58-0·69] | 1·17x10^-26^ | ** |
| Abnormal platelet count | 2·58 | [2·28-2·92] | 8·14x10^-52^ | ** | 0·62 | [0·57-0·69] | 1·97x10^-22^ | ** |
| Acellular urinary casts | 5·38 | [4·43-6·54] | 1·31x10^-65^ | ** | 0·62 | [0·5-0·76] | 6·91x10^-6^ | ** |
| Cylindruria | 5·38 | [4·43-6·54] | 1·31x10^-65^ | ** | 0·62 | [0·5-0·76] | 6·91x10^-6^ | ** |
| Hyaline casts | 5·38 | [4·43-6·54] | 1·31x10^-65^ | ** | 0·62 | [0·5-0·76] | 6·91x10^-6^ | ** |
| Abnormality of metabolism/homeostasis | 2·64 | [1·67-4·18] | 2·66x10^-5^ | ** | 0·61 | [0·49-0·74] | 1·48x10^-6^ | ** |
| Hypophosphatemia | 4·07 | [3·43-4·84] | 4·93x10^-59^ | ** | 0·61 | [0·51-0·73] | 3·90x10^-8^ | ** |
| Abnormal cardiac biomarker test | 1·43 | [1·15-1·78] | 1·24x10^-3^ | ** | 0·6 | [0·5-0·73] | 7·04x10^-8^ | ** |
| Abnormal cardiac test | 1·43 | [1·15-1·78] | 1·24x10^-3^ | ** | 0·6 | [0·5-0·73] | 7·04x10^-8^ | ** |
| Abnormality of the cardiovascular system | 1·43 | [1·15-1·78] | 1·24x10^-3^ | ** | 0·6 | [0·5-0·73] | 7·04x10^-8^ | ** |
| Decreased serum bicarbonate concentration | 2·81 | [2·34-3·36] | 7·39x10^-30^ | ** | 0·6 | [0·5-0·71] | 9·17x10^-9^ | ** |
| Increased troponin I level in blood | 1·43 | [1·15-1·78] | 1·24x10^-3^ | ** | 0·6 | [0·5-0·73] | 7·04x10^-8^ | ** |
| Increased blood urea nitrogen | 2·08 | [1·84-2·35] | 2·72x10^-32^ | ** | 0·59 | [0·54-0·64] | 4·26x10^-31^ | ** |
| Abnormal albumin level | 10·58 | [9·02-12·41] | 9·99x10^-188^ | ** | 0·58 | [0·48-0·7] | 4·27x10^-9^ | ** |
| Hyperbilirubinemia | 2·3 | [1·95-2·7] | 7·90x10^-24^ | ** | 0·58 | [0·5-0·67] | 1·14x10^-12^ | ** |
| Abnormality of coagulation | 3·29 | [2·88-3·75] | 7·24x10^-72^ | ** | 0·57 | [0·5-0·64] | 3·17x10^-22^ | ** |
| Abnormality of the coagulation cascade | 3·29 | [2·88-3·75] | 7·24x10^-72^ | ** | 0·57 | [0·5-0·64] | 3·17x10^-22^ | ** |
| Hypoalbuminemia | 10·39 | [8·85-12·21] | 7·33x10^-183^ | ** | 0·57 | [0·48-0·69] | 3·32x10^-9^ | ** |
| Abnormality of nucleobase metabolism | 6·88 | [5·62-8·44] | 1·49x10^-78^ | ** | 0·56 | [0·44-0·71] | 1·52x10^-6^ | ** |
| Abnormality of purine metabolism | 6·88 | [5·62-8·44] | 1·49x10^-78^ | ** | 0·56 | [0·44-0·71] | 1·52x10^-6^ | ** |
| Abnormality of prothrombin | 3·32 | [2·91-3·79] | 1·67x10^-72^ | ** | 0·55 | [0·49-0·62] | 4·35x10^-23^ | ** |
| Monocytopenia | 4·3 | [3·6-5·15] | 6·13x10^-59^ | ** | 0·55 | [0·45-0·67] | 1·99x10^-9^ | ** |
| Prolonged prothrombin time | 3·32 | [2·91-3·79] | 1·67x10^-72^ | ** | 0·55 | [0·49-0·62] | 4·35x10^-23^ | ** |
| Thrombocytopenia | 2·03 | [1·78-2·31] | 1·85x10^-27^ | ** | 0·53 | [0·48-0·59] | 7·09x10^-31^ | ** |
| Increased total bilirubin | 2·28 | [1·91-2·72] | 1·72x10^-20^ | ** | 0·52 | [0·44-0·62] | 4·60x10^-14^ | ** |
